# Ketosis Prevents Abdominal Aortic Aneurysm Rupture Through CCR2 Downregulation and Enhanced MMP Balance

**DOI:** 10.1101/2023.02.21.529460

**Authors:** Sergio Sastriques-Dunlop, Santiago Elizondo-Benedetto, Batool Arif, Rodrigo Meade, Mohamed S. Zaghloul, Sean J. English, Yongjian Liu, Mohamed A. Zayed

**Author notes:** These authors contributed equally to this work (see author contributions) (Sergio Sastriques-Dunlop, Santiago Elizondo-Benedetto).

## Abstract

Abdominal aortic aneurysms (AAAs) are common in aging populations, and AAA rupture is associated with high morbidity and mortality. There is currently no effective medical preventative therapy for AAAs to avoid rupture. It is known that the monocyte chemoattractant protein (MCP-1) / C-C chemokine receptor type 2 (CCR2) axis critically regulates AAA tissue inflammation, matrix-metalloproteinase (MMP) production, and in turn extracellular matrix (ECM) stability. However, therapeutic modulation of the CCR2 axis for AAA disease has so far not been accomplished. Since ketone bodies (KBs) are known to trigger repair mechanisms in response to vascular tissue inflammation, we evaluated whether systemic *in vivo* ketosis can impact CCR2 signaling, and therefore impact AAA expansion and rupture. To evaluate this, male Sprague-Dawley rats underwent surgical AAA formation using porcine pancreatic elastase (PPE), and received daily β-aminopropionitrile (BAPN) to promote AAA rupture. Animals with formed AAAs received either a standard diet (SD), ketogenic diet (KD), or exogenous KB supplements (EKB). Animals that received KD and EKB reached a state of ketosis, and had significantly reduced AAA expansion and incidence of rupture. Ketosis also led to significantly reduced CCR2, inflammatory cytokine content, and infiltrating macrophages in AAA tissue. Additionally, animals in ketosis had improved balance in aortic wall matrix-metalloproteinase (MMP), reduced extracellular matrix (ECM) degradation, and higher aortic media Collagen content. This study demonstrates that ketosis plays an important therapeutic role in AAA pathobiology, and provides the impetus for future studies investigating the role of ketosis as a preventative strategy for individuals with AAAs.

## Introduction

Abdominal aortic aneurysm (AAA) formation and rupture results from a complex series of biomolecular processes^1,2^. AAAs often remain asymptomatic during expantion, and until they spontaneously rupture, leading to a high risk of morbidity and mortality^3,4^. Unfortunately, there is currently no effective medical therapy to alleviate AAA expansion and the eventual risk of rupture. Invasive surgery is the only available management strategy for AAAs that meet traditional expansion rates and maximum aortic diameter criteria^5^. Given the limited medical treatment options for individuals with small AAAs that do not meet these criteria for operative repair, expectant management is usually the only option provided^6^. Therefore, medical stabilization of small AAAs remains an area of great interest, as this can have a substantiative impact on the longer-term management of individuals with AAA disease^7^.

Inflammation play a critical role in AAA disease progression^8^. From AAA initiation to the occurrence of microdissections, and subsequent expansion and rupture, the release of inflammatory mediators within the aortic wall is though to potentiate various molecular signals including the activation of matrix metalloproteinases (MMPs). Activated MMPs consequentially lead to extracellular matrix degradation^9–12^, and futher AAA expansion. The C-C chemokine receptor type 2 (CCR2) is an important mediator of this process by way of leukocyte trafficking to the site of aortic tissue inflammation following initial injury^13^. Previous studies have demonstrated that CCR2 knockdown can lead to attenuated AAA expansion, and our team previously demonstrated that positron emission tomography (PET) imaging of AAAs with a selective CCR2-targeted radiotracer demonstrates a high correlation with incidence of AAA rupture^14,15^. These prior studies demonstrate that CCR2 plays an important regulatory role in AAA pathogenesis, and that modulating its pro-inflammatory signaling may help alleviate disease progression.

Ketogenesis was recently discovered to impact anti-inflammatory signaling, and promote vascular tissue repair^16,17,18^. As a physiological process leading to the production of ketone bodies (KBs) such as acetoacetate (AcAc), beta-hydroxybutyrate (βHB) and acetone, ketogenesis not only serves as alternative fuel source, but also activates signaling cascades that can impact cell and organ function. Although high fat diets are linked to increased AAA expansion and aortic plaque formation^19,20^, recent studies suggested that a high fat – low carbohydrate ketogenic diet, as well as exogenous ketone body supplementation, can potentially reduce tissue inflammation and ameliorate the risk of vascular injury and atheroprogression^21,22^. It is unknown whether these potential benefits are limited to atherosclerosis, or whether ketosis can have a broader impact on degenerative aortopathies such as AAA expansion and rupture. Therefore, we hypothesized that nutritional ketosis, either in the form of a ketogenic diet or exogenous ketone body supplementation, can impact CCR2-mediated inflammation and improve MMP balance in aortic tissue to reduce the risk of AAA progression, aortic wall inflammation, extracellular matrix (ECM) content, and the incidence of AAA rupture.

## Results

### Ketosis Attenuates AAA Formation and Content of MMP9 in Aortic Tissue

Compared to animals fed a standard diet (SD), animals maintained on a ketogenic diet prior to AAA induction (KDp; **Figure 1A**) achieved a state of sustained ketosis from day 0-14 (**Figure 1B**). KDp also caused a moderate decrease in weight gain by week 1 and 2 (p < 0.001; **Figure 1C**). By week 2 there was a substantial 42% decrease in AAA diameter in KDp animals (p = 0.008; **Figure 1D and Figure S1A**). Aortic wall media demonstrated equivalent masson trichrome (MT)-stained collagen between animals maintained on SD and KDp (**Figure 1E through 1G**). Zymography analysis of harvested aortic tissue at week 2 demonstrated a modest decrease in pro and total-MMP9 in KDp animals (**Figure 1H through 1J**). These data suggest that diet-induced ketosis can inhibit AAA expansion, and that this may in part be due to a decrease in aortic wall total and pro-MMP9.

**Figure 1.**
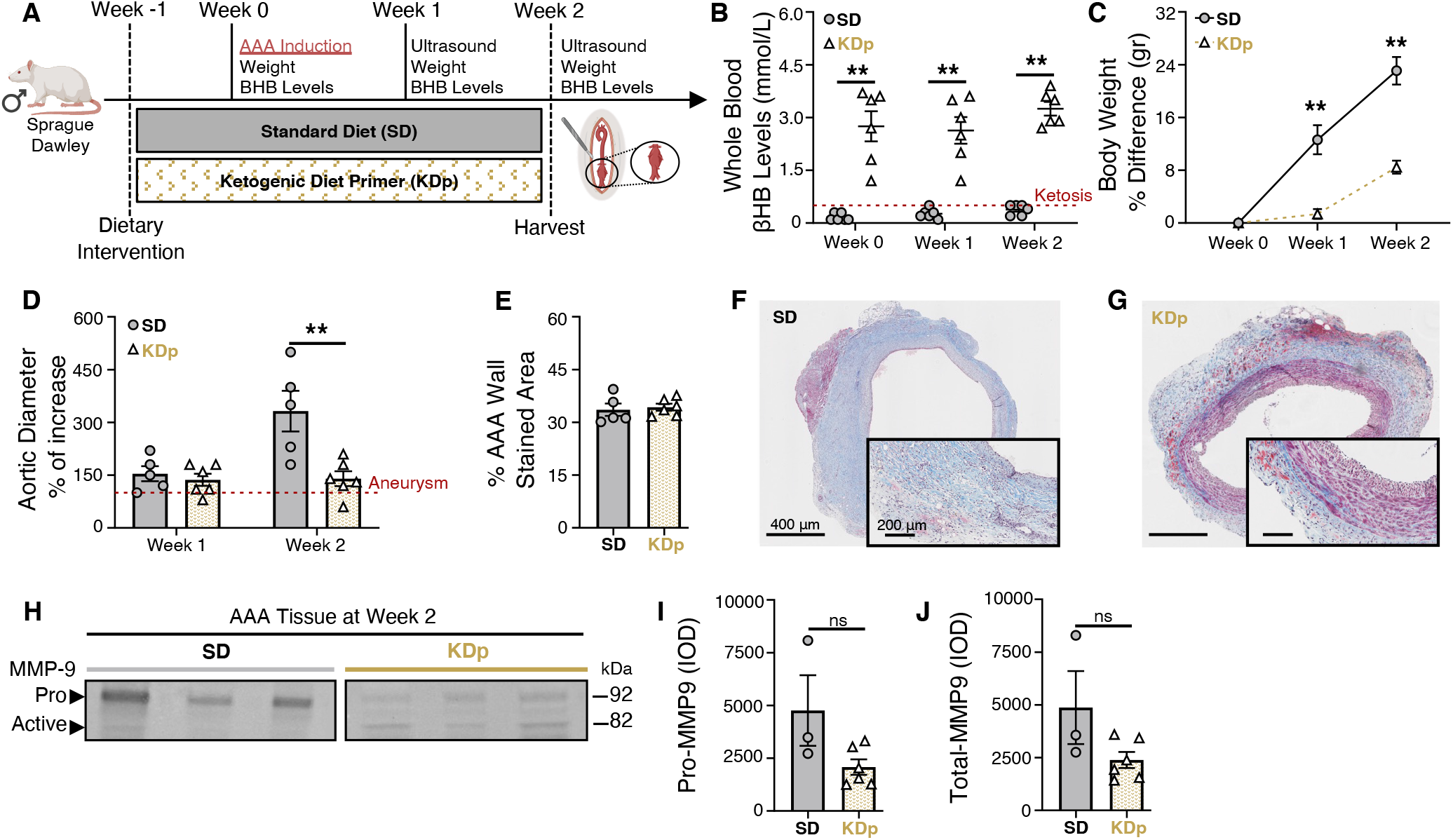
Ketosis attenuates AAA formation and MMP9. (**A**) Animals underwent exposure to PPE for AAA creation. The experimental group was given a ketogenic diet that started one-week prior to PPE exposure (KDp; N=6) while the control group was fed a standard chow diet (SD; N=5). (**B**) Ketosis (βHB whole blood levels > 0.5 mM/L) was verified at week 0, 1 and 2 in SD (0.2±0.1, 0.3±0.1 and 0.4±0.1) and KDp animals (3±1, 3±1 and 3±0.5) respectively (p < 0.01). (**C**) Percent body weight difference in SD vs KDp animals at week 1 (13±5 vs 2±1.3) and at week 2 (23±5 vs 8±2) respectively (p < 0.001). (**D**) Percent aortic diameter in SD vs KDp animals at week 1 (154±48 vs 137±42; p = ns) and at week 2 (332±129 vs 140±152; p = 0.008) respectively (aneurysms were defined by a >100% increase in the aortic diameter compared with pretreatment measurements). (**E**) AAA collagen staining quantification in percent of stained aortic tissue area for SD and KDp at week 2 (33±4 vs 34±2; p = ns) respectively. (**F** and **G**) Trichrome staining of abdominal aortas (cross-section of tissue slides) with 5x magnification for SD and KDp animals. Areas with blue staining signify areas with higher collagen deposition. (**H**) Zymogram demonstrating pro and active MMP9 levels were measured by integrated optical density (IOD).. (**I**) Pro MMP-9 levels for SD and KDp at week 2 (4.8±3×10^3^ vs 2±0.9×10^3^; p = ns) respectively. (**J**) Total MMP-9 levels for SD and KDp at week 2 (4.8±3×10^3^ vs 2.4±0.9×10^3^; p = ns) respectively. Data presented as mean ± standard deviation. ns > 0.05, *p < 0.05, **p < 0.01, ***p < 0.001 using Student’s t test. No outliers were observed in the analyses, and all data was included in the figure.

### Sustained Ketosis Reduces AAA Expansion and CCR2 Content in Rupture-Prone Animals

A sperate cohort of animals was evaluated with SD and KDp after AAA induction with PPE, and daily BAPN administration to promote AAA rupture (**Figure 2A**)^23^. By day 14, animals that survived (non-ruptured AAA; NRAAA) were evaluated via laparotomy. Animals that sustained AAA rupture (RAAA) were evaluated via necropsy. *In vivo* evaluations of aortic diameter with ultrasound (US) and positron emission tomography (PET) / computed tomography (CT) with^64^Cu-DOTA-ECL1i (selective CCR2-targeting PET radiotracer) were performed (**Figure 2A**). Animals fed KDp, and received daily BAPN, remained in a state of ketosis days 0-14 (**Figure 2B** and **Figure S2B**), and weight gain was similarly reduced at weeks 1 and 2 (p < 0.001 and p = 0.006, respectively; **Figure 2C** and **Figure S2C**). Administration of BAPN did not significantly alter βHB levels between SD and KDp animals (**Figure S3A**). Interestingly, KDp animals also had significantly reduced AAA rupture (67% vs 12%; p = 0.03; **Figure 2D and E**). Aneurysm diameter at week 1 was significantly decreased in KDp animals (p = 0.002 and p=0.01; **Figure 2F** and **Figure S1B**), and by week 2, AAA diameter was equivalent between groups. Additionally, PET/CT demonstrated significantly reduced CCR2 content in AAAs of KDp animals at week 1 (p = 0.05) and week 2 (p = <0.0001; **Figure 2G and 2H**).^18^ F-fluorodeoxyglucose PET/CT performed in KDp and SD animals at week 1 revealed comparable AAA signal uptake (p = ns; Figure S4A and S4B)^15,23,24^. These findings demonstrate that KDp animals developed smaller aneurysms with a combined 54% absolute risk reduction, and decreased CCR2 content in AAA tissue.

**Figure 2.**
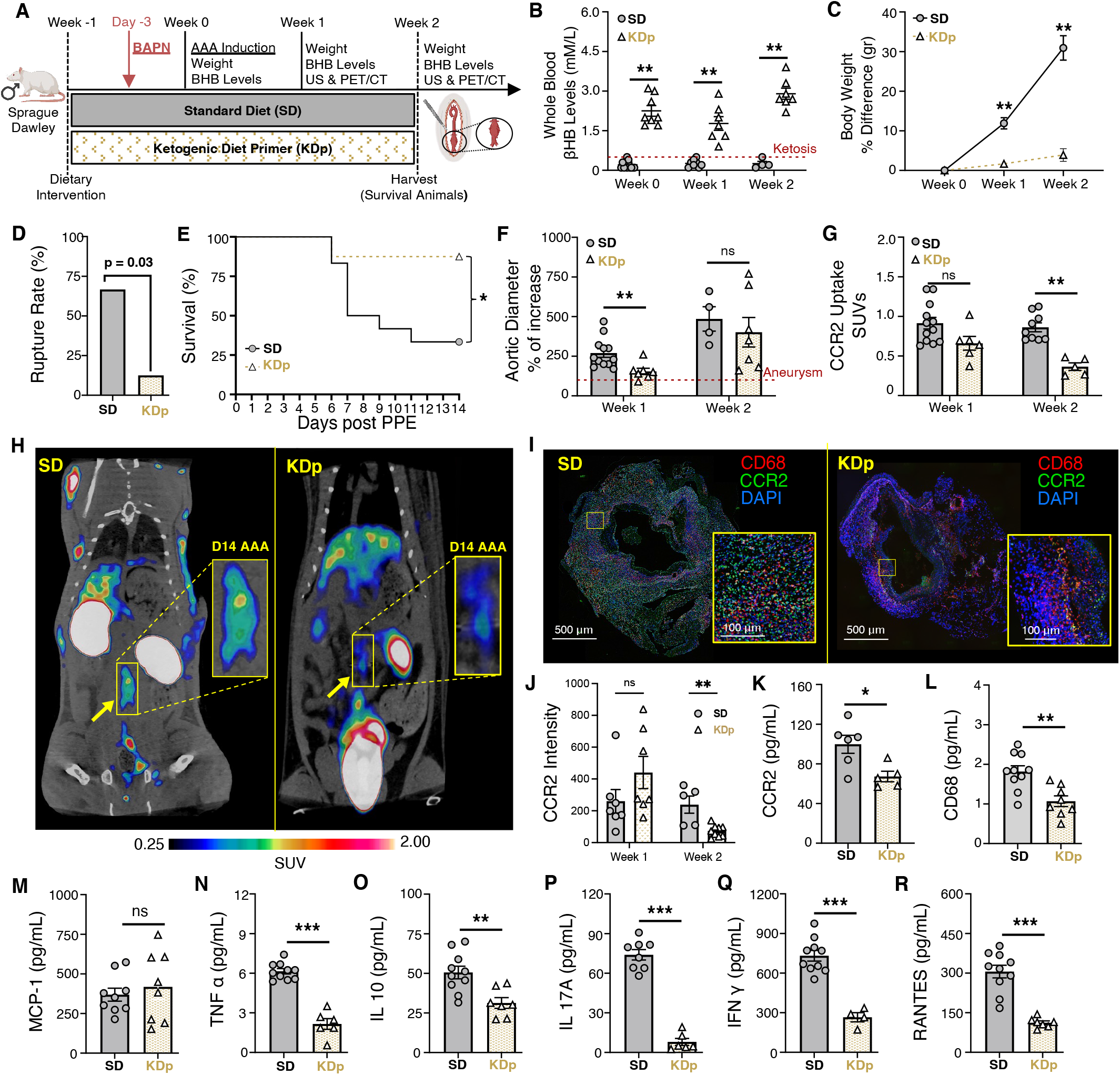
Sustained ketosis reduces AAA expansion and risk of rupture via CCR2 downregulation and Collagen 1 preservation. (**A**) Animals underwent exposure to PPE to develop AAAs and were also treated with β-aminoprionitrile (BAPN) to promote AAA rupture. (**B**) Ketosis (βHB whole blood levels > 0.5 mM/L) in SD and KDp animals at week 0 (0.2±0.1 vs 2±0.5), week 1 (0.3±0.1 vs 1.8±0.7) and week 2 (0.2±0.1 vs 3±0.5; p < 0.01). (**C**) Percent body weight difference in SD and KDp animals at week 1 (12±5 vs 2±1.3; p < 0.001) and week 2 (31±6 vs 4±3; p = 0.006). (**D**) AAA rupture event rate with statistical analysis in SD and KDp animals (p = 0.03). (**E**) Kaplan-Meier curve demonstrating rate of survival following BAPN administration. 67% (8/12) of SD animals and 12% (1/8) KDp animals developed AAA rupture. (**F**) Percent aortic diameter in SD and KDp animals at week 1 (270±93 vs 154±53; p = 0.002) and week 2 (485±153 vs 401±246; p = ns). (G) Quantitative tracer uptake of CCR2 content in AAA tissue for SD and KDp animals at week 1 (0.9±0.2 vs 0.7±0.2; p = 0.05) and week 2 (0.9±0.2 vs 0.4±0.1; p < 0.001. (**H**) Representative PET/CT coronal images at day 14 post PPE exposure showed specific and intensive detection of AAA (yellow rectangle) in SD, compared with the low trace accumulations in the KDp group of animals. (**I**) Immunofluorescence staining of abdominal aortas (cross-sectional; 5x magnification and 10x magnification) marked with CCR2 (in green: CCR2+ cells) and CD68 marker (in red: CD68 + cells; macrophages) to visualize inflammatory cells infiltration within the AAA. (**J**) Immunofluorscent intensity of CCR2 positive cells in AAA tissue at week 1 and 2. At week 1 in AAA tissue in SD and KDp animals: (**K**) CCR2 content (100±22 vs 67±12; p = 0.02). (**L**) Macrophage marker CD68 content (1.8±0.4 vs 1±0.4; p = 0.002). (**M**) Chemokine MCP-1 content (3.7±1.2 x10^2^ vs 4.2±2.2 x10^2^; p = ns). **(N)** Pro-inflammatory marker TNFα content (6.1±0.6 vs 2.1±1; p = 0.001), **(O)** IL-10 content (5±1.3 x10^1^ vs 3.1±0.9 x10^1^; p = 0.03), (**P**) IL-17A content (7.4±1.1 x10^1^ vs 0.8±0.6 x10^1^; p < 0.001), (**Q**) IFNg content (7.3±1.3 x10^2^ vs 2.6±0.7 x10^2^; p = 0.002), and (**R**) RANTES content (3±0.7 x10^2^ vs 1.1±0.2 x10^2^; p < 0.001). Data presented as mean ± standard deviation. ns > 0.05, *p < 0.05, **p < 0.01, ***p < 0.001 using Student’s t test.

### Sustained Ketosis Inhibits Cytokine Profiles Downstream to CCR2 in AAA Tissue by Week 1

The risk of AAA rupture is thought to correlate with CCR2-mediated pro-inflammatory signaling^15,25–27^. We therefore evaluated whether sustained ketosis impacts CCR2 in AAA tissue and downstream cytokine profiles by week 1 of AAA formation. We observed that immunostaining of AAA tissue in KDp animals demonstrated a marked decrease in CCR2, and CD68 macrophages (**Figure 2I** and **2J**; **Figure S5**). Correspondingly, ELISA demonstrated that CCR2 and CD68 content was significantly decreased in KDp animals at week 1 (p = 0.02 and p = 0.002, respectively; **Figure 2K and 2L**). The CCR2 ligand, monocyte chemoattractant protein-1 (MCP-1) was unchanged between KDp and SD animals (**Fig. 2M**), but the pro-inflammatory cytokines TNFα, IL-10, IL-17A, and IFN γ were decreased in AAA tissue of KDp animals (p = 0.001, p = 0.03, p < 0.001, and p = 0.002 respectively; **Fig. 2N through 2Q**). Similarly, RANTES (the ligand for C-C motif chemokine receptor 5; CCR5) was significantly reduced in KDp animals (p < 0.001; **Fig. 2R**). These results indicates that KDp notably decreases AAA macrophage infiltration, CCR2-mediated inflammation, and reduced AAA expansion and rupture.

### Ketosis Alters AAA Collagen Content and MMP Balance By Week 1

Previous work demonstrates CCR2 is known to impact MMP balance, while decreased transforming growth factor beta (TGFβ) signaling increases the risk of AAA rupture^28^. Gelatin zymography of AAA tissue at week 1 demonstrated that although pro-MMP9 and total-MMP9 were equivalent between SD and KDp animals (**Figure 3A and 3D**), active MMP9 (known to promote AAA rupture^29^) was significantly reduced in KDp animals (p = 0.03; **Figure 3B and 3D**). Similarly, total MMP 2 (known to promote AAA expansion^30^) was reduced in KDp animals (p < 0.001; **Figure 3C and 3D**). Content of MMP9 and Tissue Inhibitor of Metalloproteinases 1 complex (MMP9/TIMP1; known to prevent MMP9 over-activation) was also significantly increased in KDp animals (p = 0.008; **Figure 3E and 3F**). Correspondingly, AAA tissue in KDp animals demonstrated equivalent levels of total MMP-9 (**Figure 3G**), and significantly reduced TIMP1 compared to SD animals (p = 0.03; **Figure 3H**). Overall, these data demonstrate that sustained ketosis with KDp decreases active MMP9 while increasing MMP9/TIMP1 stabilizing complex in AAA tissue. Finally, we also observed a significant positive correlation between active MMP9 and CCR2 content in the AAA tissue in both SD and KDp animals (p = 0.03; Figure 3I and p = 0.39; Figure 3J respectively). These findings suggest that CCR2 content in AAA tissue may be responsible for activating MMPs, and therefore resulting in a higher incidence of AAA rupture.

**Figure 3.**
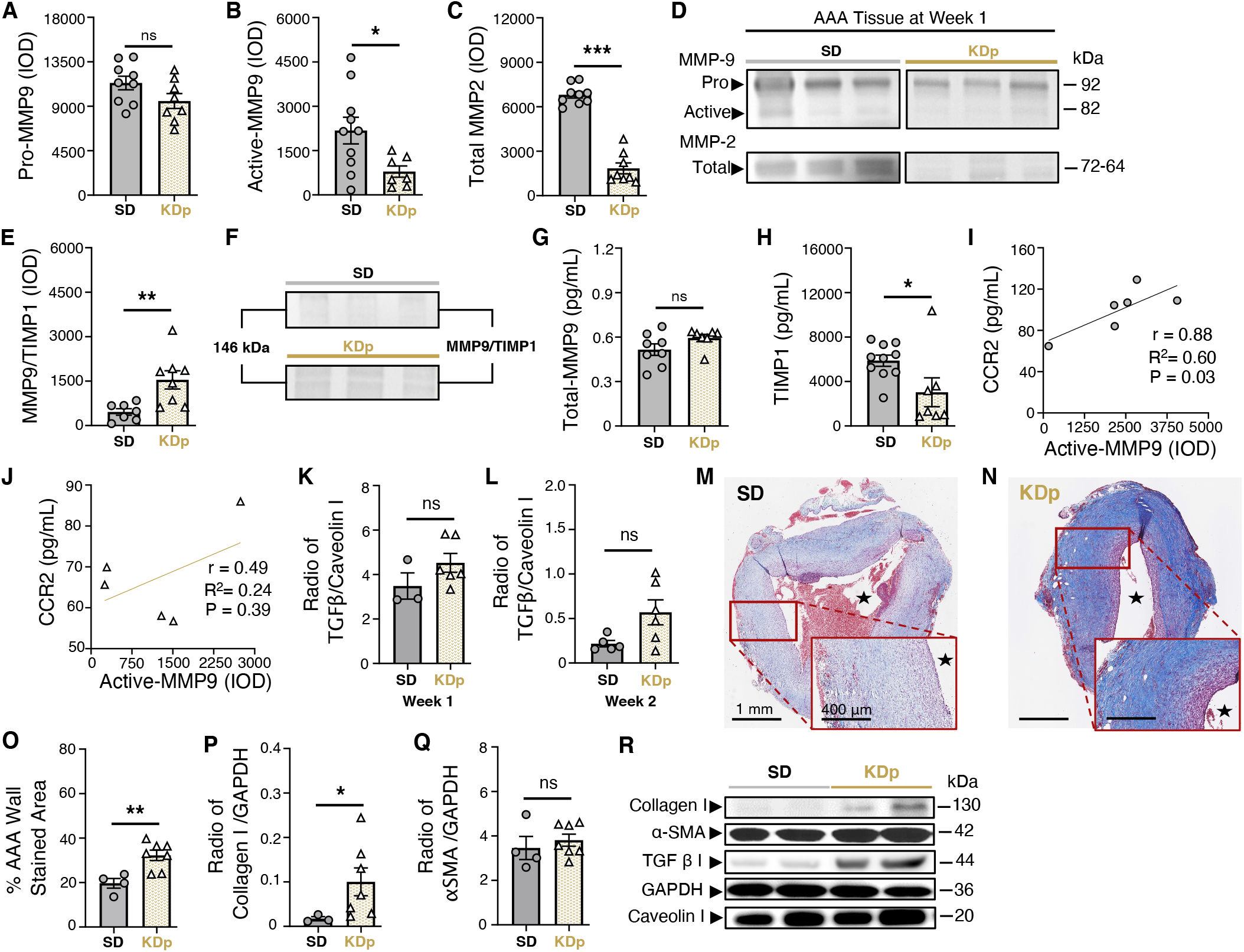
Sustained ketosis downregulates CCR2 content and inhibit its downstream signals. **(A)** Pro MMP9 levels at week 1 in AAA tissue of SD and KDp animals (11.3±2 x10^3^ vs 9.5±2.1 x10^3^; p=ns). **(B)** Active MMP9 levels in AAA tissue of SD and KDp animals (2.1±1.4 x10^3^ vs 0.8±0.5 x10^3^; p=0.03). One outlier data point in the KDp group was excluded based on pre-defined criteria prior to analysis (see methods). **(C)** Total MMP2 levels in AAA tissue of SD and KDp animals (**7**±0.7 x10^3^ vs 1.8±1 x10^3^; p<0.001). Pro and active MMP9 and total MMP2 levels were measured by integrated optical density (IOD). **(D)** Representative zymogram from AAA tissue homogenates at week 1 demonstrating pro and active MMP-9 and total MMP-2 levels in SD and KDp animals **(E & F)** Zymography demonstrating MMP9/TIMP1 complex levels in AAA tissue of SD and KDp animals (4.6±2.8 x10^2^ vs 15±8.7 x10^2^; p=0.008). One outlier data point in the KDp group was excluded based on pre-defined criteria prior to analysis. ELISA of AAA tissue homogenates in SD and KDp animals provided levels of **(G)** Total MMP9 (5±1.1 x10^-1^ vs 6±0.6 x10^-1^; p=ns), **(H)** TIMP 1 (5.9±1.6 x10^3^ vs 3.4±3 x10^3^; p=0.03). (**I** & **J**) Positive correlation analysis between CCR2 and Active MMP9 in SD and KDp respectively. (**K**) TGFβ-1 protein content expressed as a ratio to Caveolin 1 content in AAA tissue from SD and KDp animals at week 1 (2.2±0.8 x10^-1^ vs 5.7±3.4 x10^-1^; p = 0.2), and **(L)** week 2 (0.3±0.1 vs 0.4±0.1; p = 0.05). (**M and N**) Trichrome staining of abdominal aortas (cross-sectional) with 5x magnification and 10x magnification in SD and KDp animals to visualize collagen deposition in animal aortic tissue. (**O**) AAA collagen staining quantification in percent of stained aortic tissue area for SD and KDp at week 2 (20±4 vs 32±6; p = 0.006). (**P**) Collagen 1 protein content expressed as a ratio to GAPDH content in AAA tissue at week 2 for SD and KDp animals (1.4±0.8 x10^-2^ vs 10±8.2 x10^-2^, p = 0.03). One outlier data point in the SD group was excluded based on pre-defined criteria prior to analysis (see methods). (**Q)** α-SMA protein content expressed as a ratio to GAPDH content in AAA tissue of SD and KDp animals (3.4±1 vs 3.8±0.7; p = ns). (**R**) Representative Western blots of collagen 1, α-SMA, TGFβ-1, GAPDH and Caveolin 1 in AAA tissue. Data presented as mean ± standard deviation. ns > 0.05, *p < 0.05, **p < 0.01, ***p < 0.001 using Student’s t test.

Comparted to week 1, TGFβ content in AAA tissue in KDp animals trended higher in week 2 (p = ns and p = 0.05, respectively; **Figure 3K and 3L**). However, MT-staining demonstrated significantly higher collagen deposition in the AAA wall media by week 2 (p = 0.006; **Figure 3M through 3O**). In particular, type 1 Collagen content was increased in KDp animals compared to SD at week 2 (p = 0.03), while α-smooth muscle actin (αSMA) content remained unchanged (**Figure 3P, 3Q, and 3R** respectively).

### Impact of Ketosis That is Initiated ‘Therapeutically’ After AAA Formation

Animals treated with an abbreviated course of KD, therapeutically initiated 3 days post-AAA formation with PPE (KDt; **Figure 4A**), also led to a state of ketosis (**Figure 4B**). Animals treated with supplemental exogenous ketone bodies by oral daily gavage (EKB; **Figure S2** and **Figure 4A**) also led to ketosis, but only for 8-hour per day (**Figure 4B**). Similar to KDp animals, KDt and EKB animals also had reduced weight gain at both week 1 and 2 when compared to SD animals (p < 0.001; **Figure 4C** and **Figure S2**). Although, AAA rupture rate was reduced in KDt and EKB animals compared to SD animals (22% reduction with KDt, p = 0.03, and 40% reduction EKB, p = 0.12: **Figure 4D and 4E**), the relative decrease in rupture was not as much as KDp animals (**Figure 2E and 2F**). AAA absolute diameter and percentage of aortic diameter increase were also significantly reduced at both week 1 and 2 in EKB animals while only significantly reduced at week 2 in KDt animals (**Figure 4F** and **Figure S1**). These findings demonstrate that KDt and EKB therapeutic regimens lead to reduced AAA expansion and risk of rupture.

**Figure 4.**
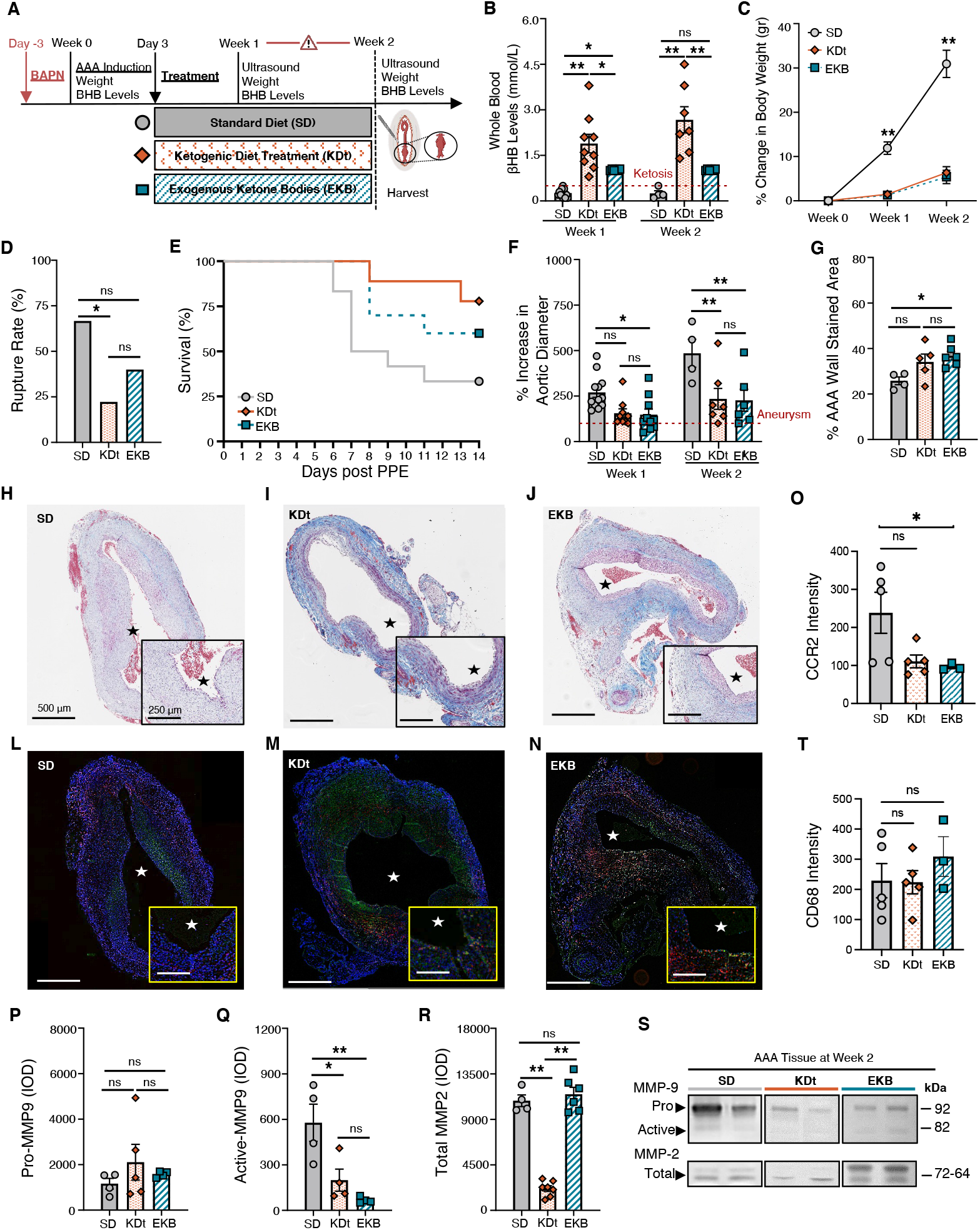
Impact of therapeutic ketosis on AAA risk of rupture. **(A)** Animals underwent exposure to PPE to develop AAAs and were also treated with BAPN to promote AAA rupture. Following AAA induction, animals received a ketogenic ‘treatment’ via an oral diet (KDt) or exogenous supplement (EKB). **(B)** Ketosis (βHB whole blood levels > 0.5 mM/L) in SD, KDt, and EKB animals at week 1 (0.2±0.1, 1.8±0.9 and 1±0.02, respectively; p < 0.05) and at week 2 (0.2±0.1, 2.7±1.1, and 1±0.02 respectively; p < 0.01) analyzed using two-way ANOVA. **(C)** Percent body weight difference for SD, KDt, and EKB animals at week 1 (11±4, 2.5±1.4, and 2.4±1.3, respectively; p < 0.01) and at week 2 (31±6, 6±3.5, and 5±3.6, respectively; p < 0.01) analyzed using two-way ANOVA. **(D)** AAA rupture event rate in SD, KDt and EKB animals (p<0.05 between SD and KDt) using survival analysis. **(E)** Kaplan-Meier curve demonstrating survival following BAPN administration. 67% (8/12) of SD animals, 22% (2/9) of KDt animals (p=0.03), and 40% (4/10) of EKB animals (p = ns) developed AAA rupture by week 2. **(F)** Percent aortic diameter at week 1 in SD vs KDt animals (270±94 and 155±73; p = 0.06), and SD vs EKB animals (148±94; p = 0.04). At week 2, in SD vs KDt animals (485±153 and 234±151; p < 0.01), and SD vs EKB animals (227±147; p < 0.01 analyzed using two-way ANOVA. **(G-J)** Trichrome staining of AAA tissue at 5x and 10x magnifications to demonstrate collagen deposition in SD vs KDt animals (26±3 and 34±8; p = ns), and SD vs EKB animals (37±5; p = 0.02) analyzed using one-way ANOVA. (**K**) CCR2 ELISA content for SD vs KDt animals (5.7±4 and 4.6±3; p = ns), and SD vs EKB animals (4.7±3; p = ns). **(L**) Pro MMP9 levels for SD vs KDt animals (1.2±0.5 x10^3^ and 2.1±1.7 x10^3^; p = ns), and SD vs EKB animals (1.6±0.12 x10^3^; p = ns). **(M)** Active MMP9 levels for SD vs KDt (5.8±2.4 x10^2^ and 2±1.4 x10^2^; p = 0.02), and SD vs EKB animals (0.7±0.2 x10^2^; p = 0.005). **(N)** Total MMP-2 levels for SD vs KDt animals (10.8±1.1 x10^3^ and 2.1±0.7 x10^3^; p < 0.001), and SD vs EKB animals (11.5±1.7 x10^3^; p = ns). Pro and active MMP9 and total MMP2 levels were measured by integrated optical density (IOD) in AAA tissue, and analyzed using one-way ANOVA. (**N**) Representative zymogram of AAA tissue homogenates at week 2, demonstrating pro and active MMP-9 and total MMP-2 levels in animals fed SD, KDt, and EKB. Data presented as mean ± standard deviation. ns > 0.05, *p < 0.05, **p < 0.01, ***p < 0.001 using one-way ANOVA, or two-way ANOVA with multiple comparison.

KDt and EKB animals also demonstrated increased AAA wall media Collagen content (p = 0.08 and p = 0.02, respectively; **Figure 4G through 4J**), and reduced CCR2 immunostaining (p = 0.06 and p < 0.05, respectively; **Figure 4O through 4N**). No difference was observed in CD68 immunostaining across groups (**Figure 4T**). Equivalent levels of pro-MMP9 were observed among both treatment groups (**Figure 4P**), but active MMP9 was significantly decreased in KDt and EKB animals (p = 0.02 and p = 0.001, respectively; **Fig. 4Q**). Total MMP2 was also notably attenuated in KDt animals (p < 0.001), but not in EKB animals (**Fig. 4R**). These data suggest that even an abbreviated therapeutic course of ketosis following AAA formation can help stabilize AAAs, preserve aortic wall collagen content, reduce CCR2 tissue content, and promote MMP balance.

## Discussion

To our knowledge our study is the first to demonstrate that diet-induced ketosis can significantly impact AAA progression and the risk of rupture. Using previously validated, pre-clinical rodent AAA models, and different ketogenic supplementation strategies, we provide a robust and comprehensive assessment of the impact of dietary ketosis on AAA formation and risk of rupture. We specifically demonstrate that administration of either a ketogenic diet (KDp or KDt) or an oral ketone body supplementation (EKB) can reliably induce systemic ketosis, significantly reduced aortic wall CCR2 and pro-inflammatory cytokines, increase collagen content in the AAA media, and promote MMP balance (Figure 5).

**Figure 5.**
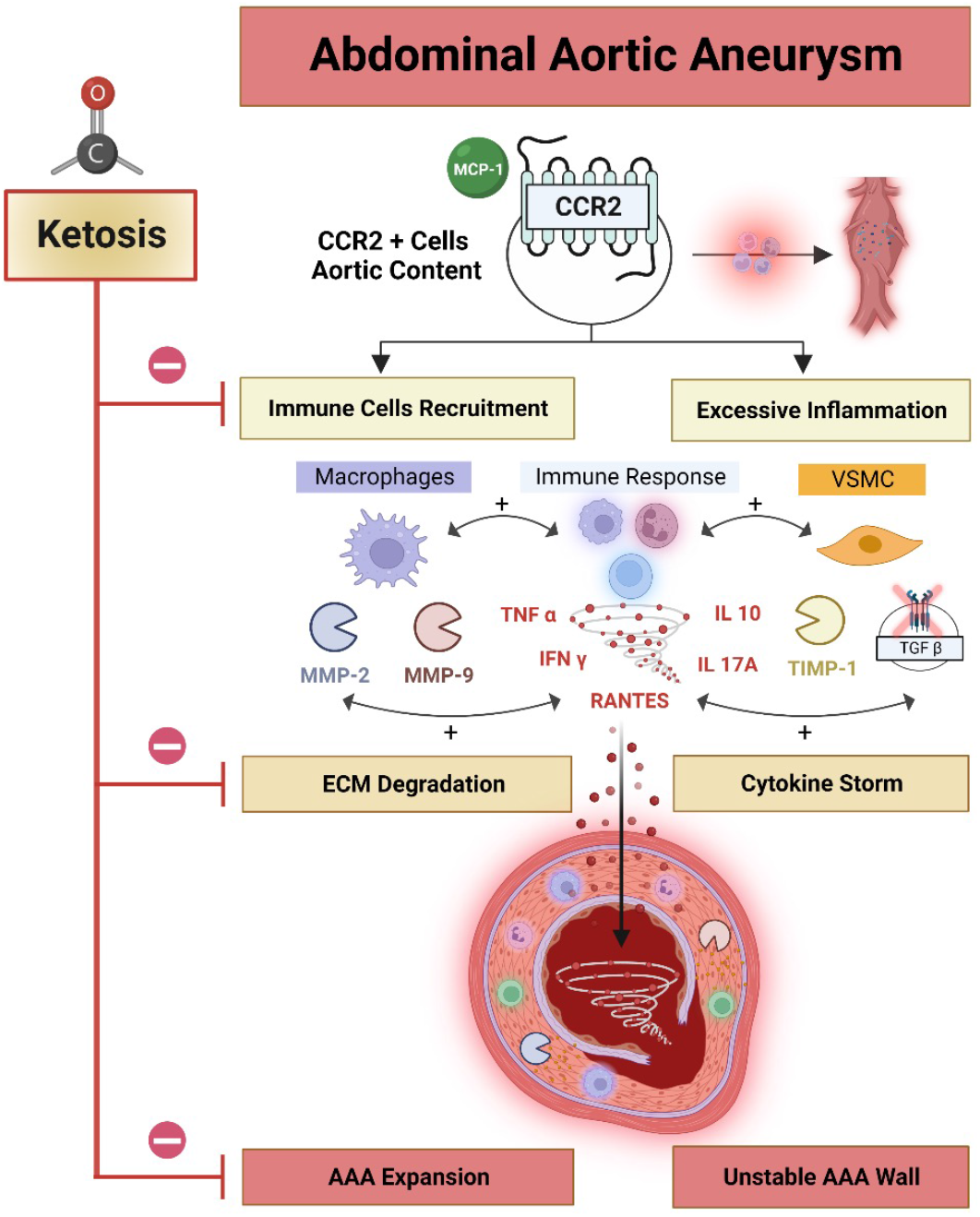
Ketosis impacts AAA expansion and risk of rupture. AAA expansion and risk rupture is influenced by CCR2, which recruits CD68+ pro-inflammatory macrophages, and also leads cytokine release, and MMP activation. Vascular smooth muscle cells (VSMCs) production of TIMP1 can complex with MMP9 to help balance out rate of MMP-medicated ECM degradation. Decreased TIMP1/MMP9 complex can lead to higher ECM degradation and AAA expansion. CCR2-mediated release of TNFa, RANTES, IL-10, IL-17A, and IFNg can further influence AAA tissue stability. Inhibition of TGFß can also lead to AAA instability and increased risk of rupture. Ketosis inhibits inflammation and ECM degradation thereby stabilizing AAA tissue and reducing the risk and incidence of rupture. Figure was made using BioRender.com.

Animals that received KDp demonstrated the most notable decrease in AAA expansion and risk of rupture. Animals that received KDt and EKB supplements also demonstrated differences in AAA progression, but not to the same extent. There was also mild to moderate variability in the of KDt and EKB on CCR2, CD68, and MMP content in AAA tissue. Administration of BAPN was reliable in inducing AAA rupture and did not appear to confound the impact of ketosis on AAA expansion and risk of rupture. Additionally, our complementary studies demonstrated that ketosis can impact pro-inflammatory CCR2-mediated signaling mechanisms that can lead to AAA progression. Therefore, this pre-clinical study demonstrates that a low-risk, and relatively easy dietary intervention, can potentially alter the course of AAA disease progression, and provides important insights that can be easily translated to human patients with AAAs who lack an effective medical management strategy.

Endogenous ketone body production mainly occurs in the liver, and results in a high glucagon/insulin ratio and increased serum free fatty acids production in the circulation^31^. This naturally occurs during periods of fasting, where βHB is released into the bloodstream as a byproduct of enzymatic degradation of ketone bodies within the mitochondrial matrix and is converted into ATP through oxidative phosphorylation^32^. βHB rises to a few hundred micromolar (μM) concentrations within 12–16 hours of fasting, 1–2 mM after 2 days of fasting^33^, and 6–8 mM with prolonged starvation^34^. Ketogenic diets modify a host’s systemic energy metabolism to mimic the biochemical impact of starvation by significantly increasing serum βHB levels, lowering blood glucose, and increasing fatty acid concentrations^35^. These regimens were originally introduced as a treatment for refractory epilepsy in children and have now become popular for weight loss programs, patients with diabetes, obesity, various types of cancer, and among high performance athletes^36–40^. Standard ketogenic diets that are devoid of carbohydrates can lead to elevated βHB serum levels that are consistently >2 mM^40^. Recent studies demonstrate that βHB can serve as an important signaling mediator that can inhibit histone deacetylases^41^, blunt tissue oxidative stress^42,43^, active G-protein-coupled receptors^44,45^, and regulate inflammatory mediators such as prostaglandin D2^46^, interleukins^47^, nuclear factor kappa B (NF-κB)^48^, and NLRP3 inflammasome^49^. Similarly, our study shows that animals with high serum βHB have blunted tissue inflammation and CCR2 content, which in part likely contributes to reduced pathological AAA expansion and risk of rupture.

Uniquely, our study administered three different ketosis regimens: two types of ketogenic diets (KDp and KDt), and an oral supplement regimen (EKB). KDp included a 1-week priming period prior to AAA formation, that imitates the phenomenon of keto-adaptation that occurs in humans who are maintained longer-term on a ketogenic diet^50^. This regimen aided in determining whether a ketosis primer can have a ‘protective’ impact against AAA formation and expansion. On the other hand, KDt was designed to evaluate the potential ‘therapeutic’ impact of ketosis on expansion and rupture of AAA post-induction with PPE. This regimen would hypothetically be similar to how medical management would be prescribed in humans with small AAAs that do not yet meet the traditional size criteria for operative intervention. In the course of this study, we observed that animals tolerated both KDp and KDt, and that both were successful in inducing a sustained systemic state of ketosis. Interestingly, both regimens yielded significant reductions in AAA expansion and incidence of rupture relative to animals that received SD. However, the longer-term KDp regimen appeared to have a more protective impact, and a more impressive reduction of CCR2 content in AAA tissue. These findings suggest that the length of diet-induced ketosis may be an important variable in the extent of reduction of AAA tissue inflammation and risk of rupture.

With the recent advent of EKB supplements, oral regimens have been increasingly utilized to manipulate levels of circulating blood ketone body concentrations in humans for various health benefits^51^. While most studies involving EKB supplementation have traditionally focused on its impact among high-performance athletes^52^, these supplements are increasingly being studied as remedies for conditions such as epilepsy, heart failure, diabetes, and sepsis-related muscle atrophy^53^. Our study evaluated the use of EKB to induce ketosis in animals with AAAs that are prone to rupture. Interestingly, we observed that EKB not only decreased AAA tissue inflammation (Figure S7), but also reduced AAA expansion and incidence of rupture (**Figure 4**). The impact of EKB on CCR2 content and AAA rupture was variable from KDp and KDt, and we suspect this is because EKB only induced intermittent ketosis (limited to 8 hours per day). Nonetheless, these findings are the first to show that oral supplementation with ketone bodies can indeed serve as a minimally invasive method for the potential medical management of AAAs, and is a compelling topic for further exploration in future human clinical trials that completement prior efforts^55–58^.

Our study results also suggest that ketosis has a multifaceted impact on aortic wall structure and function. Inflammation is the major molecular mediator of AAA disease progression (**Figure 5**). Previous studies demonstrated that excessive aortic wall inflammation can inhibit reparative signaling, wall fibrosis, and collagen deposition, which can in turn accelerate AAA expansion and lead to a higher risk of rupture^59^. Tissue macrophages are known to promote AAA disease, in particular subsets that highly express CCR2^12^. We as well as others, also previously demonstrated that genetic or molecular targeting of CCR2 can reduce AAA progression^13–15^. Here we provide further compleing evidence that CCR2 content indeed correlates with AAA disease progression, and that systemic ketosis *in vivo* can significantly reduce its both CCR2 content as well as downstream pro-inflammatory cytokines in AAA tissue.

Previous studies investigating the inflammasome in AAA tissue, demonstrated that TNFa and RANTES are both up-regulated in expanding AAA wall tissue^60,61^. Inhibition of TNFa appears to decrease aortic wall MMP activity, reduce ECM disruption, and decrease aortic diameter in a murine pre-clinical AAA model^62^. In another study, Empagliflozin, a sodium-glucose cotransporter 2 inhibitor that increases plasma ketone bodies^63,64^, was found to reduce aortic aneurysm diameter and aortic wall RANTES in Apo E -/- mice^65^. Similarly, in our study we observed that diet-induced ketosis can significantly decrease aortic wall pro-inflammatory cytokines TNFa and RANTES, as well as increase aortic wall Collagen content. Although the direct mechanism of action for this is yet to be fully elucidated, we suspect that the molecular interplay between macrophage and other pro-inflammatory cell types may be playing a critical role in the immune modulation of these processes and AAA progression^66,67^

A central pathological feature of AAA disease progression is excessive and aberrant extracellular matrix (ECM) remodeling. This results from increased MMP activity, which promotes rapid ECM breakdown and disruption of the integrity of the aortic wall^68,69^. Previous work demonstrates that MMP2 plays a central role in the formation and early expansion of AAAs, while MMP9 is more related to late AAA expansion and risk of aneurysm rupture^23,70,71^. Synergistic activation of both MMP2 and MMP9 provides an unfavorable environment that can accelerate AAA dilation and lead to a higher risk of aneurysm rupture^72^. Previous studies also demonstrate that ketosis, high serum βHB, and signaling via NF-Kβ, play key roles in suppresses MMP-9 expression in colonic tissue^73^. Our studies extend on this molecular mechanism of action, and demonstrate that ketosis and elevated serum βHB can also significantly attenuate both active MMP9, and total MMP2 in aortic tissue. In fact, a CCR2 antagonist has shown to downregulate MMP-9 expression in lung cancer cells, therefore mitigating cellular motility and metastatic invasion^74^. These results may help explain why we observed a notable decrease in MMP-9 content in AAA tissue from animals with ketosis.

TIMPs are endogenous specific inhibitors of MMPs produced by vascular smooth muscle cells (VSMCs) as well as other cell types in AAA tissue^75^, which inhibit zymogenesis of pro-MMPs and reduces overall MMP activation. Given their central role in maintaining the dynamic balance in ECM turnover in aortic wall tissue, the role of TIMPs in AAA progression continues to be an area of intense investigation^29^. Our study demonstrates that while nutritional ketosis decreases the content of free TIMP1, it significantly increases the content of the stabilizing TIMP1/MMP9 complex in AAA tissue. This data suggests that complexed TIMP1 leads to a reduction in active MMP9 content, therefore decreasing AAA wall ECM degradation, further aneurysm expansion, and the overall risk of rupture (**Figure 5**).

Our study also demonstrated a mild-moderate, but non-significant, increase in AAA tissue TGFβ content in animals treated with ketogenic diets (**Figure 5**). TGFβ belongs to a superfamily of growth factors that regulate many cellular functions such as cell growth, adhesion, migration, differentiation, and apoptosis^76^. TGFβ content appears to be significantly reduced in human AAA tissue^77^. A recent study demonstrated that ketosis promoted TGFβ-induced myocardial fibrosis and Collagen 1 and 3 deposition in spontaneously hypertensive rats^78^, suggesting that TGFβ upregulation was deleterious in this setting. However, in aortic tissue, TGFβ appears to have a beneficial role. For example, administration of TGFβ neutralizing antibodies appeared to promote excessive monocyte-macrophage infiltration within murine and rat AAA tissue^28,79^, while overexpression or administration of TGFβ1 significantly increased aortic wall collagen deposition^80^, and collagen synthesis in normal arteries^81^. This in part explains our observation that animals receiving a ketogenic diet had significantly increase aortic wall Collagen 1 deposition, which correlated with higher aortic tissue TGFβ content.

We acknowledge that there are some limitations in our study. First, all our data is derived from pre-clinical rodent models that are not necessarily representative of human AAA pathophysiology. However, the rat AAA rupture model was previously validated and shown to be the most reliable and consistent AAA rupture model currently available. Second, our studies did not systematically evaluate arterial blood pressure. This would have required sophisticated in dwelling sensors and the use of continuous telemetry. While such monitoring systems are feasible for shorter experimental protocols, our 2–3-week experimental protocol would have greatly complicated the experimental design and led to several confounding variables. We therefore elected to instead serially monitor AAA endpoints via ultrasound, which provided reliable and reproducible data. Third, our study used a single composition for the ketogenic diet intervention. We acknowledge that this is not fully representative of the wide variety of lipid and oil-based ketogenic diets consumed by humans, but this was selected to maintain consistency and adherence within all rodent study groups.

In conclusion, this study demonstrates that a ketogenic diet and EKB supplementation strategy that can significantly reduce AAA expansion and reduce the incidence of AAA rupture. Importantly, a ketogenic priming period appears to also be further protective, while EKB appears to be less effective than other dietary regimens. Ketogenic diets reduced CCR2 content, promoted MMP balance, and attenuated ECM degradation in AAA tissue. These findings provide the impetus for future pre-clinical and clinical studies geared to determine the role of ketosis as a medical management tool for human patients with AAAs that do not yet meet the criteria for surgical intervention.

## Methods

### Animals

Male Sprague-Dawley rats (200–300g) were obtained from Charles River Laboratories (Wilmington, MA) and used for all described experiments. All animals were housed at 21 °C in a 12/12 h light/dark cycle and had access to food and water ad libitum. Anesthesia was administered with a mixture of ~1.5% isoflurane and oxygen for all procedures. The core body temperature was monitored and maintained with a heating pad (37°C). Use of all animal models was approved by the Institutional Animal Care and Use Committee (IACUC) at Washington University School of Medicine in St. Louis.

### Induction of AAA

Rats were induced to develop infrarenal AAAs via an established model using porcine pancreatic elastase (PPE; 12 U/mL) as previously described^24^. Ventral abdominal wall laparotomy is performed, and the infrarenal abdominal aorta was exposed. (**Figure S8**). A customized polyethylene catheter (Braintree Scientific, Braintree, MA) was introduced through an infrarenal aortotomy, and elastase was infused into the isolated aortic segment for 30 minutes. The exposed aortic segment was dilated to a maximal diameter, and constant pressure was maintained with the use of a syringe pump. Using a video micrometer, the baseline maximum aortic diameter was measured. After 14 days, all rat aortas were re-exposed via ventral abdominal laparotomy, maximal aortic diameters were measured, and aortic tissue was harvested for further analysis (**Figure S8**).

### Promoting AAA Rupture

As previously described, β aminopropionitrile (BAPN) is reported to promote AAA tissue inflammation by day 6, and AAA rupture between days 7 and 14, but unlikely to cause rupture after day 14 if it has not already occurred^15^. We therefore promoted AAA rupture with daily administration of BAPN on a specific cohort of rats (RAAA) starting 3 days before PPE exposure. Through drinking water, 300mg BAPN was administered daily (0.3% BAPN in 25mL water consumed per day by a 250g rat)^82^. AAA growth was monitored for 1 week (6-7 days) or 2 weeks (14 days). At the 1 or 2-week timepoints, rats were sacrificed, AAA diameters were evaluated, and aortic tissue was harvested for further analysis. Rats that developed ruptured AAAs during the study period promptly underwent necropsy to confirm and analyze the pathology (**Figure S8**). Tissue harvested at week 1, prior to rupture, was mainly used to assess AAA tissue inflammation (**Figure S2**).

### Animal Diets

We evaluated four different dietary interventions. First, control groups in the AAA (n=5) and RAAA (n=12) cohorts were fed with a standard chow diet (SD). Second, experimental groups in the AAA (n=6) and RAAA (n=8) cohorts were fed a very high fat diet with almost no carbohydrate, also known as a classic ketogenic diet one week prior to PPE exposure to induce a ‘priming’ keto-adapted status^83^ before AAA induction (**Figure S9**). Ketogenic diet was then maintained prospectively in these groups following AAA formation. Third, experimental groups in the RAAA (n=9) cohorts were separately started on ketogenic diets as a ‘treatment’ intervention 3 days after AAA induction. Lastly, experimental RAAA rats were administered a SD along with exogenous ketone body (EKB) supplementation starting 3 days after PPE exposure: RAAA + EKB (n=10). As previously described^84^, this EKB supplementation was performed with daily intragastric gavage of 1,3-Butanediol (BD; 5g per kg dose; Prod # B84785-100ML, St. Louis, MO) and animals achieved a ketosis state only for 8 hours per day (**Figure S9**).

### CCR2 PET/CT Scan Evaluation

#### Synthesis and Radiolabeling of DOTA-ECL1i

The ECL1i peptide (LGTFLKC) was synthesized from D-form amino acids by CPC Scientific (Sunnyvale, CA). DOTA-ECL1i was prepared following our previous report. Copper-64 (64Cu, t1/2=12.7 hour) was produced by the Washington University Cyclotron Facility. The DOTA-ECL1i conjugate was radiolabeled with 64CuCl2 (64Cu-DOTA-ECL1) as described, and the radiochemical purity was determined by radio-HPLC^15^.

#### Animal PET/CT Imaging and Image Analysis

Dynamic PET scan and corresponding CT images were obtained using Inveon MM PET/CT (Siemens, Malvern, PA) at 45 to 60 minutes after a tail vein injection of 64Cu-DOTA-ECL1i (11.1 MBq per rat) to minimize the effect of blood retention on AAA uptake. To localize tracer uptake, a CT contrast agent (1.0 mL, eXIA 160XL, Binitio, Canada) was administrated via tail vein after PET imaging. Contrast-enhanced CT (Bin of 2, 90 mm axial FOV, 60 kV, 500 μA, 500 ms exposure time, 10 ms settle time, no magnification, pixel size: 80–100 μm) was performed. The AAA uptake was calculated as standardized uptake value (SUV) in 3-dimensional regions of interest from PET images without correction for partial volume effect using Inveon Research Workplace software (Siemens) (30). Dynamic (0–90 minutes) 18F-fluorodeoxyglucose (41.1 MBq per rat) PET was also performed in AAA rats at week 1 and 2 post PPE exposure. Only a specific number of each group of rats received PET/CT imaging and analysis.

#### Ultrasound Aortic Assessments

Noninvasive ultrasound (GE, 12 MHz Zonare, Mountain View, CA), was used to evaluate serial aortic maximum diameter measurements. Relative to baseline aortic diameter prior to PPE exposure, the percentage increase in aortic diameter was evaluated at 1- and 2-weeks post-PPE exposure. As previously described, aortic aneurysms were defined as >100% increase in the aortic maximum diameter relative to baseline diameter^24,85^.

#### Blood βHB Assessments

Animal state of ketosis was evaluated via whole blood D-βHB (Keto-MoJo blood ketone meter; Keto-Mojo, Napa, CA, USA) concentrations^86^. Tail vein puncture was used for blood sample, which was tested on day 0 pre-PPE exposure, and then 1- and 2-weeks following AAA induction. βHB values > 0.5 mmol/L were indicative of ketosis.

#### Animal Weight

Animal whole body weights were evaluated at day 0 pre-PPE exposure, and 1 and 2 weeks followed AAA induction. Body weight was evaluated by the difference between the values at the baseline (Day 0) and the values at week 1 and 2 respectively and then divided by the baseline to assess difference. All these absolute numbers were then multiplied by 100 to present it as the percentage of difference in weight throughout the time of the study.

#### Histology and Immunostaining

Aortic tissue was harvested from all animals. AAA tissue was fixed in Histochoice (VWR), and paraffin embedded. Paraffin blocks were sectioned at 5 μm, and deparaffinized. Processing for antigen retrieval was performed with Sodium Citrate solution, pH 6.0, for 10 min. Tissue sections were blocked with 10% serum, and sections were incubated with primary antibody anti-CD68, 1:100 [Bio-Rad, MCA341GA]. Sections were then incubated with anti-mouse secondary antibodies conjugated with HRP (Cell Signaling), DAB peroxidase substrate kit (Vector Laboratories), and counter stained with hematoxylin, imaged using an Olympus fluorescent microscope system. To evaluate AAA tissue morphology and pathology, tissue sections were also evaluated using Hematoxylin and Eosin (H&E) and Mason Trichrome (MT). AAA wall tissue-stained sections were then analyzed and quantified by Image J software via color deconvolution and shown as percentage of stained area for specific regions of interest (ROI).

#### ELISA and Cytokine Array Assessments

AAA tissue protein was extracted using RIPA buffer with proteinase inhibitor (Sigma #MCL1). Protein quantification was done by Bradford assay. For each AAA tissue samples 25ug of protein was analyzed for Tissue Inhibitor of Metalloproteinases 1 (TIMP1)-specific ELISA (RayBiotech, ELR-TIMP1-CL-2), MMP-9 (MyBioSource, MBS722532), CD68 (MyBioSource, MBS705029) and Cytokine multiplex assay (Millipore, RECYTMAG-65K) using manufacturer instructions.

#### MMP Activity Zymography

For each AAA tissue sample, 25ug of protein was loaded on wells of 10% Gelatin Zymogram electrophoresis gels. Gels were then incubated in Zymogram renature buffer for 30 min, followed by 36 hours of Zymogram development buffer at 37°C. Gels were then stained with Coomassie Brilliant Blue R-25 solution from BioRad for 30 min, followed by destaining buffer (20% Methanol, 20% Acetic acid, 60% DI water) until MMP bands were visualized. Gels were scanned on BioRad Chemi doc and analyzed using ImageJ software.

#### Immunoblotting

From each AAA sample 25μg of denatured protein was separated on 4-12% polyacrylamide by electrophoresis and then transferred to PVDF membrane. The membranes were then incubated with collagen type-1 (Millipore #ABT123, 1:2000), TGF-β1 (Sigma #AV44268, 1:1000), and α-SMA (Sigma #A2547, 1:1000). GAPDH (Sigma # G9545, .1mg/ml) and Caveolin-1 (Santa Cruz # sc-53564, 1:1000) were used as loading controls. Membranes were treated with HRP-conjugated secondary antibody at room temperature for 1 hour and evaluated with chemiluminescence. Blot band intensities were analyzed using ImageJ software.

#### Statistical Analysis

All data are presented as the mean ± SD. Most group comparisons were performed using unpaired t test. For comparisons that included one endpoint in more than one animal/diet groups, an ordinary one-way ANOVA with multiple comparison was performed. For comparisons that included more than one endpoint in more than one animal/diet group, we utilized a two-way ANOVA with multiple comparison. Data was considered statistically significant with p ≤ 0.05. Kaplan-Meier curve was generated to assess the survival of BAPN-exposed animals. GraphPad Prism 9 (La Jolla, CA) was used for all statistical analyses and graphical data representations. In certain circumstances outlier data points were excluded from the analysis if they met the pre-determinated criteria of the outlier was more than (1.5 x Interquartile Range (IQR)) above the third quartile (QR) or below the first quartile (Q1). MT cross section staining’s were analyzed using ImageJ software by color deconvolution, adjust threshold and region of interest assessment of the AAA wall.

## Non-standard Abbreviations and Acronyms

AcAc: Acetoacetate
BAPN: β-aminopropionitrile
BD: 1,3-Butanediol
CCR2: C-C chemokine receptor type 2
ECM: Extracellular Matrix
EKB: Exogenous ketone bodies
H&E: Hematoxylin and Eosin
IACUC: Institutional Animal Care and Use Committee
KB: Ketone Bodies
KD: Ketogenic Diet
MMP: Matrix Metalloproteinase
MT: Masson trichrome
NF-κB: Nuclear factor kappa B
PPE: Porcine Pancreatic Elastase
QR: Quartile
RAAA: Ruptured AAA
ROI: regions of interest
SD: Standard Diet
TIMP1: Tissue Inhibitor of Metalloproteinases 1
VSMCs: Vascular smooth muscle cells
αSMA: α-smooth muscle actin
βHB: beta-hydroxybutyrate

## Acknowledgments

- Conceptualization: Sergio Sastriques-Dunlop, Santiago Elizondo-Benedetto, Sean J. English, Mohamed A. Zayed
- Methodology: Batool Arif, Sergio Sastriques-Dunlop, Santiago Elizondo-Benedetto, Rodrigo Meade, Mohamed S. Zaghloul
- Investigation: Sergio Sastriques-Dunlop, Santiago Elizondo-Benedetto, Batool Arif
- Data Collection: Santiago Elizondo-Benedetto
- Supervision: Sean J. English, Mohamed A. Zayed
- Writing—original draft: Santiago Elizondo-Benedetto, Mohamed A. Zayed
- Writing—review & editing: Santiago Elizondo-Benedetto, Sergio Sastriques-Dunlop, Mohamed A. Zayed

## Sources of Funding

This work was supported by grants from National Institute of Health, National Heart Lung and Blood Institute R01HL153436 (Mohamed A. Zayed), R01HL150891 (Mohamed A. Zayed), R01HL153262 (Mohamed A. Zayed).

## Disclosures

The authors declare that they have no competing interests.

## Data and Materials Availability

All data are available in the main text or the supplementary materials.

## Supplemental Material

Supplemental Results; Supplemental Methods; Figures S1 to S9

## Notes

### Competing Interest Statement

The authors have declared no competing interest.

